# PERCEPTIVE: an R Shiny pipeline for the prediction of epigenetic modulators in novel species

**DOI:** 10.1101/2024.11.20.624555

**Authors:** Eric M. Small, Christina R. Steadman

## Abstract

Epigenetic processes play key roles in regulating gene expression, genome stability, and metabolic output in organisms across the tree of life. Yet, the role epigenetics plays in regulating genomes and behaviors remains underdeveloped for microalgae, particularly as new species are identified and characterized. This is likely due to the cumbersome nature and species-dependent attributes of epigenetic wet-lab methodologies, which preclude the rapid identification of epigenetic modifications and modulators. However, there is extraordinary conservation of epigenetic processes from budding yeast to humans; in many cases, one may infer how behavior and function are epigenetically regulated in novel species by simply identifying epigenetic modulators, or the proteins responsible for conferring epigenetic modifications. To this end, we have developed a graphical software package, titled PERCEPTIVE (<u>p</u>ipelin<u>e</u> for the predi<u>c</u>tion of <u>ep</u>igenetic modulators in no<u>v</u>el sp<u>e</u>cies). This novel platform solely uses the genomic sequence of an organism, and preexisting information from model species, to predict the epigenetic modulators and associated modifications in a novel species. Predictions are presented to the user in a graphical interface, which provides literature-based interpretation of results, enabling users to quickly understand potential epigenetic processes in their species of interest and plan follow-up experiments. To test PERCEPTIVE, we predicted epigenetic modulators in several feedstock candidate algae species. To validate these predictions, wet-lab studies were performed, including mass spectrometry; these results underscore the high accuracy of PERCEPTIVE predictions. Overall, PERCEPTIVE represents a powerful tool for the research and manipulation of algal species, which does not require *a priori* knowledge of epigenetics and is accessible to a broad set of investigators.

## INTRODUCTION

Algae continue to provide a rich repertoire of biomolecules that can be harnessed for production of food, feed, fuels, and products. Indeed, continuous concerted efforts toward laboratory and environmental screens have identified new species with novel traits and desirable characteristics—metabolic, physiological, and phenotypic—indicative of feedstock potential [1,2]. Characterization studies, ranging from genotyping to phenomics, provide foundational information for deeper understanding of microalgae physiology and function. However, microalgae, along with other organisms, do not behave as predicted, particularly during scale-up cultivation and production, where environmental variance influences behavior and phenotype. Ensuring consistent behavior is essential to assure quality and yield to achieve bioeconomy production goals. As such, it is likely we are missing key biological information in microalgae [3].

In the genomic age, it is both relatively inexpensive and informative to sequence the genome of a novel species to gain insights into its behavior. Computational models to predict coding regions, gene products, and homology to proteins in model species transform genomic sequencing into a powerful tool for this purpose. For example, the homologous proteins for fatty acid metabolism that a species possesses can inform the conditions and nutrient requirements that may favor the generation of a desired bioproduct or precursor [4]. This ability to ascertain (with high likelihood) the proteins and processes that are active in a novel organism has fueled genetic engineering of non-model species, with the goal of improving organism behavior. Synthetic biology techniques to introduce transgenes, knock out endogenous genes, or enhance expression of genes (via gene editing) have led to improved yield and higher performance of feedstocks [5,6]. Nonetheless, outcomes are not always clear-cut; for example, the introduction of a gene may not result in the consistent production of a protein or bioproduct. This can be for many genetic reasons; however, we, and others, hypothesize that epigenetic mechanisms likely underlie many of these disparate outcomes [7,8]. For example, the incorporation of a new gene in a telomeric region that has lost methylation of the core histone H3 on lysine 79—and is natively heterochromatic (or transcriptionally repressive)—may result in poor expression of the gene [9,10]. This irregular expression may shift with epigenetic context, physiological conditions, or cell cycle. Similarly, editing of a native gene to upregulate expression may not prove effective for similar reasons. However, transgene incorporation, or gene upregulation, could potentially be rescued by manipulation via epigenetic machinery. Moreover, epigenetic machinery plays key roles in directing physiological states, ranging from the cell cycle to nutrient responses [11–13]. Indeed, previous work demonstrates that inhibition of epigenetic machinery responsible for DNA methylation profoundly increases lipid accumulation in the feedstock species *Picochlorum soleocismus* [14]. Yet, epigenetic modifications and their associated machinery—though prominent—are among the least studied and leveraged mechanisms for enhancing algae behavior [8,14].

Despite the promise of epigenetic manipulation, traditional methods to uncover potential epigenetic mechanisms are costly and time consuming wet-lab experiments, and in many cases, produce limited results [15]. We propose that a bioinformatics approach is both more resource and time efficient and is more likely to yield better downstream experimental design. While epigenetic modifications are not directly genomically encoded, and available genomes alone provide little epigenetic information, epigenetic modifications *are* conferred by genomically encoded chromatin modifying enzymes (CMEs). Additionally, this chromatin machinery has been extensively studied in model species [16,17]. While much remains to be uncovered, the highly conserved nature of CMEs and the core subunits of chromatin (histones) allow us to infer much about an novel organism’s behavior and function, particularly if homologous proteins are present in a species of interest [16]. Unfortunately, existing algal genome annotations, whether bioinformatically or experimentally defined, provide scant clues about the epigenetic processes in algae species, requiring extensive bioinformatics mining and literature review to uncover the epigenetic repertoire of a given species [18].

Based on these principles, herein, we describe a novel software package, PERCEPTIVE (<u>p</u>ipelin<u>e</u> for the predi<u>c</u>tion of <u>ep</u>igenetic modulators in no<u>v</u>el sp<u>e</u>cies) to predict the epigenetic processes in a novel organism that an end-user can employ prior to extensive wet-bench experimentation. PERCEPTIVE has four distinct goals and attributes as follows:

1. the full annotation of poorly or unannotated genomes without the necessity for command-line tools, which is accessible to an everyday computer user;
2. the identification of homologs of chromatin modifying enzymes from unassembled or assembled genomes of novel species;
3. the prediction of DNA and histone covalent modifications; and
4. the identification of ideal wet-lab reagents to study chromatin features in novel species.

Critically, this package reduces the necessity for wet-lab experiments to identify CMEs in species that may lack certain chromatin modifications and processes, or for which available lab reagents are unsuitable. For example, this may be an organism that lacks 5mC DNA methylation or an organism with significant sequence divergence in histone proteins around a residue of interest, which commercial reagents may not be able to readily detect [15,19]. Crucially, this package removes the requirements for a user to be familiar with command-line tools, as all data is analyzed and presented in a graphical user interface (GUI). Moreover, and most importantly, the tool does not require *a priori* knowledge of epigenetics and incorporates expert knowledge and a vast literature review to uncover potential epigenetic mechanisms active in a novel species for the end-user.

To validate our pipeline, we performed mass spectroscopy for histone modifications and utilized previous wet-lab observations of DNA methylation in select feedstock algal species. The predictions produced by PERCEPTIVE have high concordance with experimental observations, which reinforce the pipeline’s predictive efficacy. Finally, while our focus is predominately on algal feedstock species, epigenetic mechanisms can explain behavior and responses to environmental perturbations or conditions across nearly all species. We believe that the deeper understanding of epigenetic mechanisms in non-model species, which can be derived from this package, could improve our understanding of a vast array of disciplines, from pathogen responses to plant behavior and even ecosystems dynamics.

## METHODS

### 2.1 PERCEPTIVE pipeline design

PERCEPTIVE is composed of two software components: a Linux application that facilitates the annotation of a genome from an assembled genome or unassembled reads, and a helper application that is agnostic of the operating system (Windows, Linux, etc.), which extracts annotated CMEs from the output of the first application and lends interpretation and wet-lab analysis. The first application consists of an R shiny graphical interface that wraps several established software applications into a simple bash-like pipeline. As outlined in Figure 1, either assembled genomes in *fasta* format (which are preferable) or trimmed unassembled long or short reads in *fastq* format can be used as input to PERCEPTIVE.

**Figure 1.**
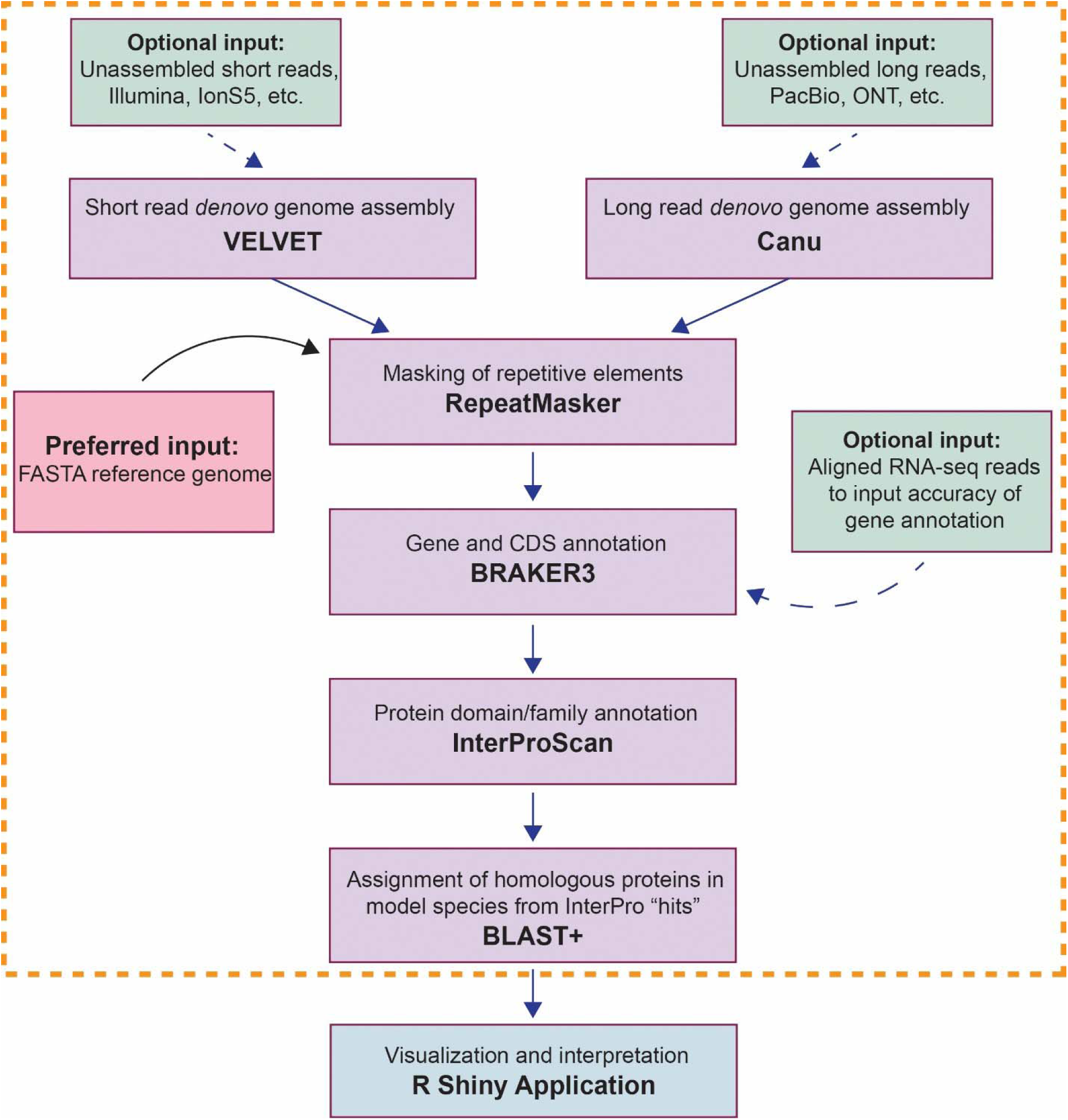
Bioinformatic Flowchart: Shown is a flowchart of the PERCEPTIVE pipeline bounded in orange, which is feed into the R Shiny PERCEPTIVE application for visualization.

If unassembled reads are provided, VELVET is used to assemble a genome from short reads (Version 1.2.10), or Canu is used to assemble a genome from long reads (Version 2.2) [20,21]. Following genome assembly, or input of assembled genomes, PERCEPTIVE employs RepeatMasker to mask tandem repeats in a clade-dependent fashion to prevent downstream spurious identification of coding regions or protein homologs (Version 4.1.7) [22]. Subsequently, masked genomes are used as an input to the well-described BRAKER pipeline to identify putative coding regions and open reading frames (Version 3.0) [23]. Optionally, RNA-seq data can be input to BRAKER to improve annotations. The resulting peptide sequences corresponding to coding regions are then used as input for InterProScan (Version 5.70-102) prediction of protein domains and families [24,25]. InterProScan was chosen for its implementation of hmmer and the InterPro consortia’s broad well-characterized collection of hidden Markov models. Following annotation of protein coding regions with domains and families, the annotations are parsed using a curated list of InterPro accessions. The protein sequences of any domain/family hits are subsequently blasted for sequence homology against the protein coding sequences of well-studied model organisms (*A. thaliana*, *C. elegans, C. reinhardtii*, *D. melanogaster, D. rerio, H. sapiens*, *M. musculus*, *S. cerevisae*, *T. thermophila, S. pombe*, *X. laevis, Z. mays*) using NCBI’s BLAST+ (Version 2.16.0) [26]. More poorly studied species were excluded from this database of model species to avoid uninformative outputs, like “uncharacterized protein of putative function,” which arise when less studied species like *Ectocarpus spp.* are included. While this approach favors specificity, it should be expected that some proteins may not yield BLAST results in highly divergent, esoteric species. While this may limit some downstream user tuned applications (e.g., literature review of a specific homologous methyltransferase), it does not limit the predictive capacity of PERCEPTIVE, which utilizes hidden Markov Models as core evidence.

The final output of this pipeline includes a repeat masked *fasta*; a Gene Transfer Format (GTF) file of all annotated coding regions; a General Feature Format 3 (GFF3) file of all InterPro family and domain annotations; BLAST+ hits for all protein sequences with InterPro annotations corresponding to a curated list of domains important to chromatin modifying enzymes; and the protein sequences of histone proteins identified by InterPro. This package of files is subsequently read into the R Shiny helper application, which will run in any Linux, OS X, or Windows environment capable of running an R package [27,28]. This application parses the annotation data to provide a literature-based interpretation and prediction of epigenetic modifications that may be present in an organism. Both applications are provided as Singularity images to avoid the complexity of setting up the pipeline environment (Version 4.2.0) [29].

### 2.2 Genome data

Assembled unmasked genomes for *Nannochloropsis gaditana, N. oceanica,* and *N. salina; Tetraselmis striata;* and *Picochlorum soleocismus* were downloaded from the Joint Genome Institute. Accessions B-31, CCMP1779 v2.0, CCMP1776, Tetstr1, and DOE101 respectively.

### 2.3 Culture conditions

*N. gaditana* was cultivated as previously described [15]. Briefly, triplicate cultures were cultivated in 125 mL shaker flasks (100 rpm) under 16/8 light:dark cycle in f/2 with 1% CO_2_ and in 200 μEm^−2^s^−1^. Cells were collected 1 hr prior to lights on and 3 hr after lights on. *Tetraselmis striata* was cultivated as previously described [30,31]. Briefly, triplicate cultures were cultivated in 2L shaker flasks (90 rpm) under 16/8 h light:dark cycle in f/2 media with 350Lppm sodium nitrate at 8.25 pH (with supplemented CO_2_ as needed). Aliquots were taken daily across the growth cycle for 10 days, until nitrogen starvation. Aliquots from both species were centrifuged, decanted, and flash frozen with liquid nitrogen until processed.

### 2.4 Histone extraction and mass spectroscopy

2x10^8^ *N. gaditana* or *T. striata* cells were resuspended in 300 μl Nuclear Isolation Buffer. (NIB contains 15 mM Tris-HCl (pH 7.5), 60 mM KCl, 15 mM NaCl, 5 mM MgCl2, 1 mM CaCl_2_, 250 mM sucrose, and 0.3 % NP-40.) 1 mM DTT, 1:100 Halt protease inhibitor, and 10 mM sodium butyrate were added and cells were lysed by eight freeze thaw cycles: 5 minutes on dry ice, followed by 5 minutes in a heat block at (37°C). Nuclei were then pelleted at 600*g* for 5 minutes at 4 °C. Nuclei were subsequently washed twice with NIB without NP-40, and re-pelleted as above. Nuclei were lysed and histones were solubilized with five volumes of 0.2 M H2SO4 for 1 hour at room temperature. Insoluble cellular debris was removed by centrifugation at 4,000 ×g for 5 minutes at 4 °C. Soluble histones were precipitated from the supernatant by addition of trichloroacetic acid to a final concentration of 20 % (v/v) for 1 hour at room temperature. Histones were pelleted at 10,000*g* at 4°C for 5 minutes. The supernatant was discarded, and the histone pellet was washed in ice-cold acetone + 0.1% HCl, and repelleted as above and allowed to air dry. Pelleted purified histones were sent to the Northwestern Proteomics Core for subsequent analysis on dry ice. Histones were derivatized and digested according to Garcia *et al* [32]. Histones were derivatized using propionic anhydride reagent for 1 hour prior to digestion with 1 µg of Promega sequencing grade trypsin overnight at 37 °C. After digestion, histone peptides were derivatized with propionic anhydride for 1 hour. Peptide samples were dried by vacuum centrifugation then re-suspended in 0.1 % trifluoroacetic acid prior to MS analysis. Targeted LC-MS/MS was performed on a Thermo Scientific TSQ Altis triple quadrupole mass spectrometer using a 70-minute method. Peptides were analyzed using selective reaction monitoring (SRM) LC-MS/MS on 3 technical replicates per sample. A blank was performed after each third replicate. Raw data were analyzed using Skyline software as detailed in previously published work [33,34]. The maximum average of three biological replicates, and two physiological conditions (Light/Dark *N. gaditana*) or two biological replicates and three physiological conditions (Nitrogen Replete, Deplete, Starved *T. striata*) were used for visualization and confirmation of post-translational modifications. A cutoff ratio of 1:99, modified to unmodified peptides was used to define a positive confirmation of a given modification.

## THEORY

### 3.1 Prediction schema

For an organism to possess a DNA or histone covalent modification, two key elements must be present. One, the presence of a modifiable substrate, and two, the presence of an enzyme capable of conferring a modification. In the case of DNA modifications, the first requirement is always true, as these modifications are conferred on two of the four nucleotide bases that make up DNA in all organisms: cytosine and adenine [35]. Additionally, for the second requirement, the enzymes responsible for DNA modifications are substrate specific, in that an enzyme that confers 5-methyl cytosine (5mC) cannot confer 6-methyl-adenine (6mA). In this case, any evidence for an enzyme which can confer a DNA modification, whether 4-methyl-cytosine (4mC), 5mC, 5-hydroxy-methyl-cytosine (5hmC), or 6mA is sufficient to predict that an organism may possess and utilize DNA modifications to regulate its DNA and transcription. In this case, the predictive capacity of PERCEPTIVE is binary, with species that are predicted to lack DNA modifying enzymes having zero probability of DNA methylation and species with DNA modifying enzymes having the highest probability of DNA methylation. Covalent histone modifications, on the other hand, are much more nuanced. While the histone proteins are highly conserved, in many species they have undergone significant duplication events leading to a number of isoforms with similar function but divergent peptide sequences [17,36]. Moreover, many of the chromatin modifying enzymes specific for histones are promiscuous and redundant in nature; for example, Gcn5 acetylates the core histone H4 on lysines K5, K8, K12, K14 and on the core histone H3 at K9 and K27, while HAT1 acetylates H4 on K5 and K12 [37,38]. Furthermore, the histone deacetylase proteins are equally promiscuous with Rbd3 (HDAC1 in humans), removing at least H4: K5ac, K8ac, and K12ac, for example [39]. Despite this promiscuity, some histone modifications are placed by specific enzymes, such as the methylation of K79 on H3 by Dot1 [10]. Adding to this complexity, histone modifications exhibit trans crosstalk, for example H3K79me and H2B ubiquitylation at K123 (K120 in metazoan), with H3K79me requiring placement of H2BK123ub prior to its deposition, and H4K16ac further driving H3K79me [40–42]. This represents three distinct problems for predicting histone modifications: 1) varied histone residue sequences and multiple copies of the histone proteins; 2) the promiscuity of histone modifying enzymes; 3) and histone modifications crosstalk and dependence. To address the degrees of certainty presented by these problems, PERCEPTIVE uses an additive scoring matrix for evidence with probability descriptions as follows: Best = 1, Excellent = 0.875, Better = 0.75, Good = 0.625, Fair = 0.5, Poor = 0.375, Negligible = 0.25, and Zero = 0. To manage the first problem, it is necessary to identify a modifiable substrate. Using HMMs for each of the core histones, we identify genes predicted to code for histones. On account of the fundamental nature of hidden Markov models, we cannot discriminate between histone isoforms, e.g., H3.0 and H3.1, which have discrete roles and modifications in humans and other metazoan. As such, PERCEPTIVE treats all histone isoforms as core histones, e.g., in the case of H3.0 and H3.1, as H3. Additionally, while subsequent analysis could provide clues about which model metazoan isoforms may correspond to those in a novel species, histone isoform modification and utilization for regulation varies between model species, making prediction complex and variable. To account for mutations and evolutionary divergence, the commonly applied Levenshtein edit distance formula is used to find the closest alignment between a novel histone protein sequence and model core histone sequence, which results in the fewest number of insertions, substitutions or deletions [43]. Then, if any protein sequence identified as a core histone retains an appropriate residue for modification in the correct position the modification is given 0.5 points (Figure 3).

**Figure 2:**
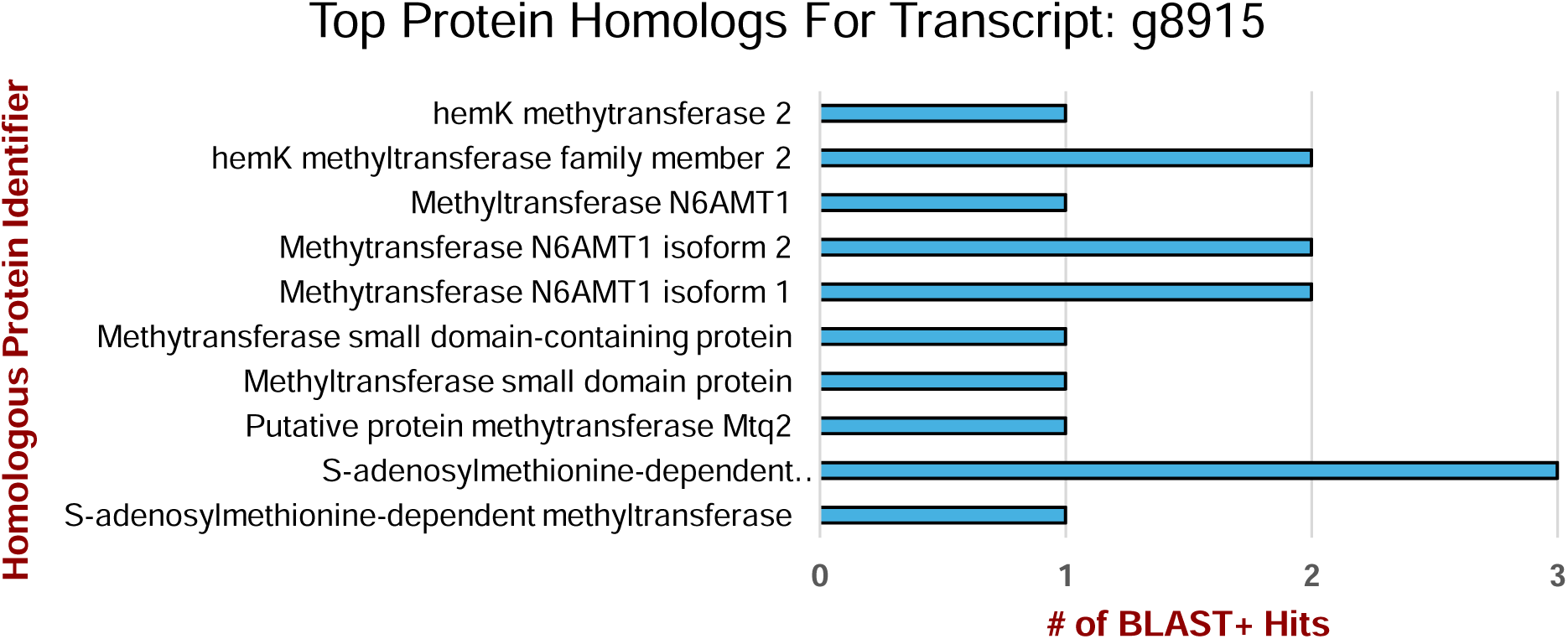
Representative graph of BLAST+ hits from several model organisms for transcript g8915, which was annotated by InterPro with evidence as a protein with DNA methylase, N-6 adenine-specific function. The subsequent viewing window provides BLAST metrics and data about model organisms in which homologous regions were found.

**Figure 3.**
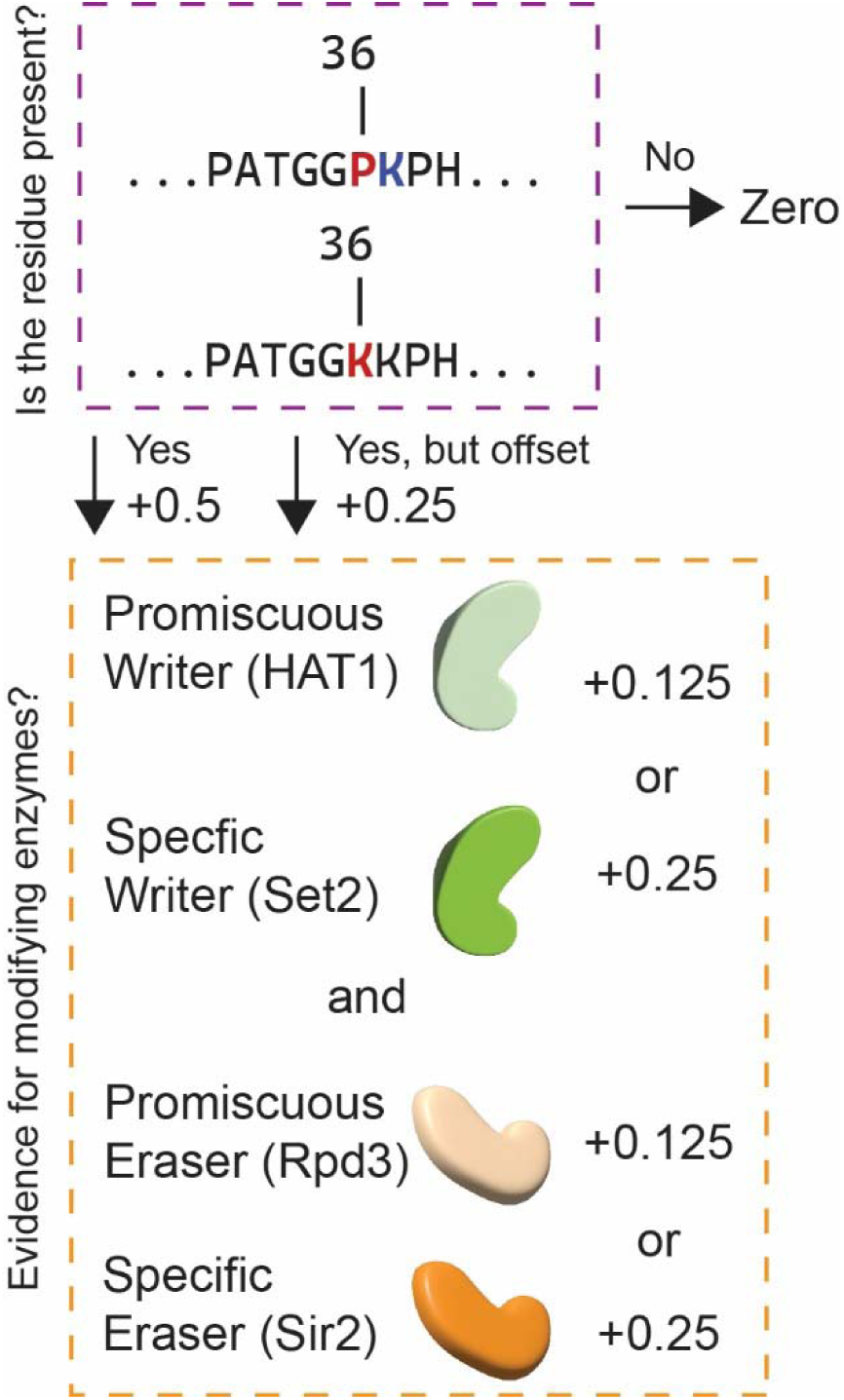
PERCEPTIVE Decision Tree: PERCEPTIVE assigns points for modifiable amino acid residues in histone proteins, and then points for evidence for chromatin modifying enzymes that may write, erase, or read modifications on that residue.

To account for the possibility of extensive insertions and deletions, PERCEPTIVE also assigns 0.25 points if an appropriate modifiable residue is found one residue up or downstream of the expected residue, for example, H3K35 instead of H3K36 (Figure 3). If no modifiable residue is found, the modification receives zero points, the probability is zero, and no other evidence is considered. To address problem two, writer and eraser chromatin modifying enzymes are scored separately: however, if any enzyme which can either confer or remove a modification is found, however promiscuous, 0.125 points is added to the score for both the presence of a writer and the presence of an eraser. Then, the specificity of an enzyme is considered, with enzymes with specific non-promiscuous functions, for example, Dot1, receiving an additional 0.125 points. (Figure 3). In the case of problem three, if evidence for a modification required for a subsequent modification to be written is not found, points for a modification are retained, but a penalty of 0.125 points is taken (e.g., evidence is found for H3K79Kme but not H2Bub). In practice, this means that if PERCEPTIVE identifies a histone H3 protein with a lysine at position 36, evidence of the specific methyltransferase Set2, and evidence for a non-specific demethylase, the modification H3K36me is given 0.5, 0.25, and 0.125 points respectively, for a total of 0.875 points, or excellent probability. Finally, it should be noted that PERCEPTIVE, with the exception of H3K9me, does not discriminate between mono-, di-, and trimethylation of a residue, as most histone modifications are conferred in a processive manner by the same CME, making it impossible to predict the degree of methylation an organism will possess (from the presence of the CME alone). That being said, in almost all cases, it should be expected that an organism with the correct CMEs should have some degree of mono-, di-, and trimethylation.

**Figure 3:**
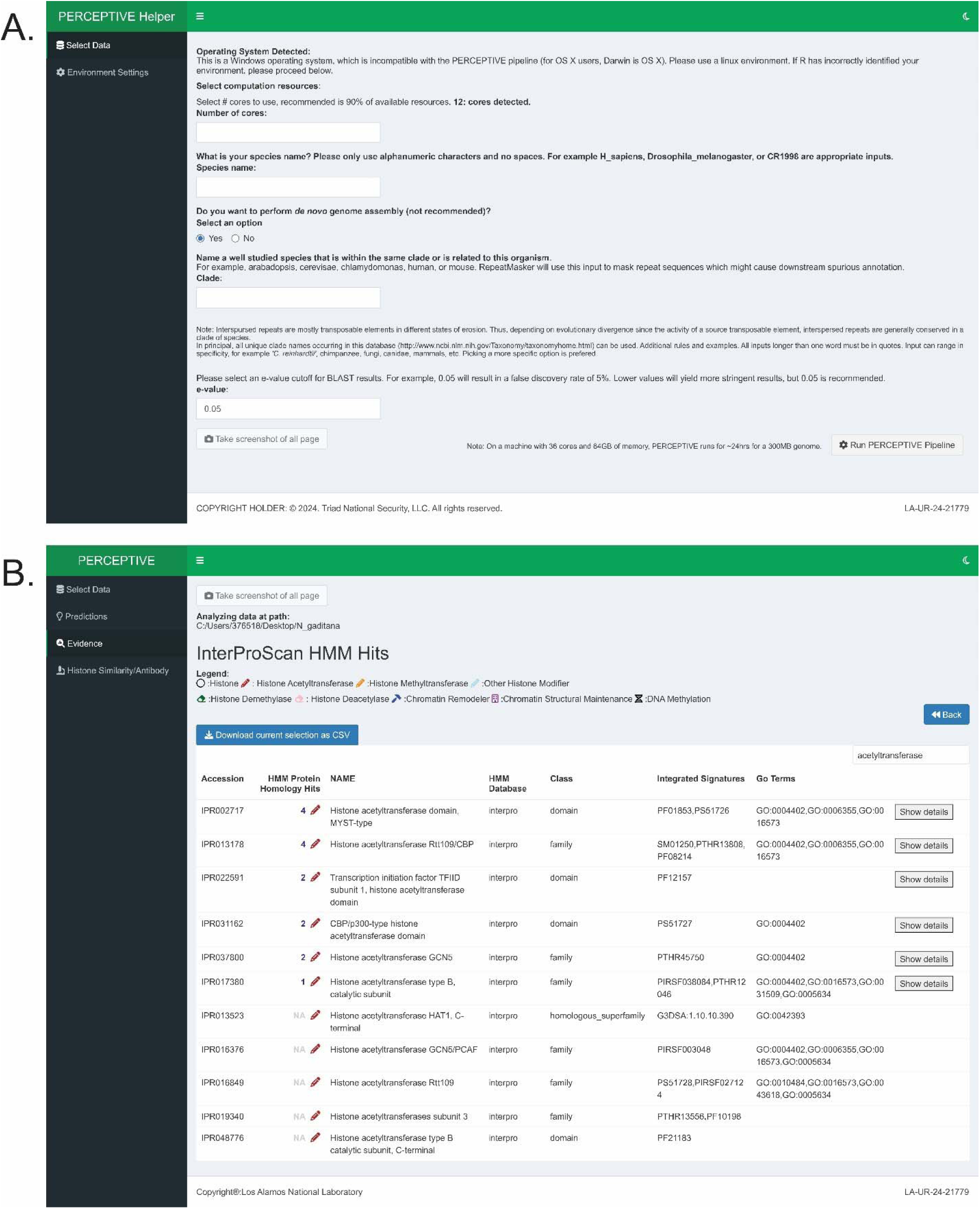
The R shiny PERCEPTIVE Interface. A) Graphical Linux-based PERCEPTIVE pipeline helper prompts users for inputs specific to their organism of interest and launches the PERCEPTIVE pipeline. B) PERCEPTIVE displays hidden Markov model evidence for histone acetyltransferases in *N. gaditana*.

In addition to reporting predictions for chromatin modifications, PERCEPTIVE also reports evidence for other classes of CMEs, namely chromatin remodeling enzymes and chromatin structural proteins. Given the variable functions of these enzymes, and the roles of these proteins in numerous structural elements, predictions for chromatin remodeling and structural elements are not provided. Nonetheless, evidence for these features is reported and can provide critical insights into the function of the chromatin machinery in a novel species.

### 3.2 Evidence for PERCEPTIVE: Hidden Markov model implementation and BLAST

Hidden Markov models (HMMs) have been extensively applied to the problem of annotating proteins in novel genomes. HMMs share some distinct advantages over local alignment search tools like BLAST, which wholly rely on sequence similarity to identify the function of a punitive coding region [44]. While HMMs are less specific than tools like BLAST, for example, HMMs poorly predict specific proteins (e.g., a specific 6mA methyltransferase like human N6AMT1), they effectively identify protein domains with annotated functions like “DNA methylase, N-6 adenine-specific.” The permissive nature of HMMs to changes in protein sequence is ideal, however, as evolutionary pressures are more likely to drive mutations in protein regions not deleterious to their function while favoring the conversation of essential domains. As such, we took advantage of the InterPro database of HMMs, trained on well-characterized species, to annotate protein coding regions with more general “domain-like” functions, like “DNA methylase, N-6 adenine-specific” to avoid missing potential punitive proteins with epigenetic functions on account of sequence divergence. Nonetheless, BLAST-like searches do offer advantages for the planning of downstream wet-lab work and literature review. As such, following InterPro annotation, protein sequences of interest, for example, a protein coding region annotated with “DNA methylase, N-6 adenine-specific,” are blasted against a database of well-studied model organism proteins. BLAST results, in this case, may return that a “DNA methylase, N-6 adenine-specific,” annotated protein, is most similar to human N6AMT1, allowing the user to further research N6AMT1 and plan wet-lab experiments, which might be enabled by commercial antibodies, etc., (Figure 2). Accordingly, this data is presented under the evidence tab in PERCEPTIVE. However, owing to the diversity of gene repertoires, varying gene functions, and divergent gene nomenclature across organisms, BLAST results are not utilized as a component of the prediction schema below, and predictions solely rely on available evidence from HMMs.

## RESULTS

### 3.1 Overview

The PERCEPTIVE pipeline (<u>p</u>ipelin<u>e</u> for the predi<u>c</u>tion of <u>ep</u>igenetic modulators in no<u>v</u>el sp<u>e</u>cies) predicts and informs the user of potential epigenetic modifications in any eukaryotic organism of interest in four key steps, including 1) annotating the genome, 2) identifying CME homologs, 3) predicting the epigenetic modifications from those identified CMEs, and 4) providing information on wet-lab experiments to do for verification. Step one is addressed by the Linux-dependent graphical pipeline, which performs assembly and annotation of *de novo* or poorly annotated genomes (Figure 3A).

Steps two through four are addressed by a graphical helper application (which is agonistic to the operating system) and presents the predicted DNA and histone modifications for an organism, including outlining the underlying evidence for the predicted modifications in addition to evidence for chromatin structural and remodeling enzymes from InterPro and BLAST+. It also visualizes sequence conservation of the core histone proteins as compared to model species, aiding the user in the planning downstream wet-lab experiments (Figure 3B). To verify the accuracy of PERCEPTIVE, we ran the genomes of several algal species with known epigenetic modifications, or modifications directly measured, through the pipeline, including *Nannochloropsis gaditana, N. oceanica, N. salina, Picochlorum soleocismus,* and *Tetraselimis striata*.

### 3.3 Validation

To validate the prediction of CMEs capable of conferring DNA methylation, we analyzed the genomes of *N. gaditana, N. oceanica, N. salina,* and *P. soleocismus*. Our previous work has shown that *Nannochloropsis spp.* lack 5mC and 5hmC methylation but do possess 6mA methylation [15,45]. Moreover, our previous studies in *P. soleocismus* have shown a robust phenotypic response to 5AZA, a drug that blocks the CME responsible for 5mC, suggesting that *P. soleocismus* possesses the epigenetic machinery to confer 5mC [14]. Moreover, a previous painstaking gene-by-gene examination of *P. soleocismus* genome annotations found evidence for DNA methylating CMEs specific to 5mC [14]. Predictions for a representative *Nannochloropsis spp.*, *N. gaditana*, and *P. soleocismus* suggest that PERCEPTIVE accurately predicts CMEs for DNA methylation (Table 1). For example, in *N. gaditana* we do not predict 5mC methylation, aligning with our previous findings in which biochemical assays, whole-genome bisulfite sequencing, and third generation Oxford Nanopore sequencing find no evidence for 5mC in *N. gaditana* [15]. The apparent strength of our predictions comes with the caveat that DNA methyltransferases are not promiscuous, and as such, predictive scores for DNA methylating CMEs are binary. In other words, either there is evidence for DNA methylation, resulting in a “best” score, or no evidence resulting in a ”zero” score.

**Table 1.**
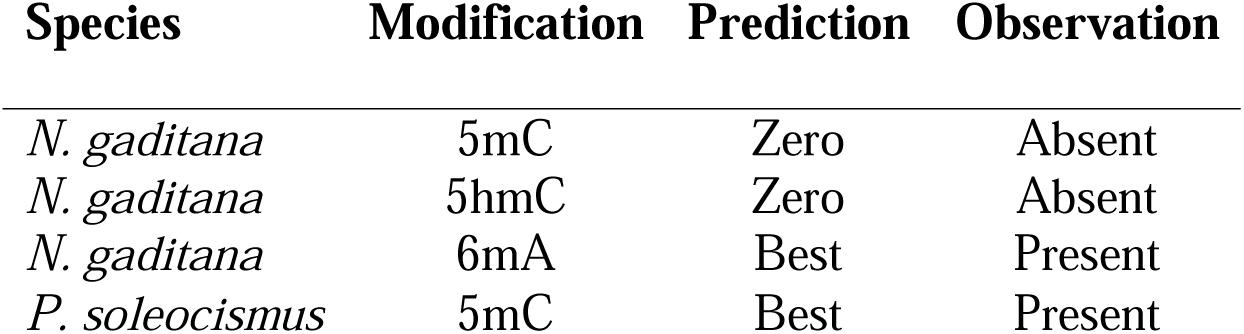
Validation of Chromatin Modifying Enzymes for different types of DNA Methylation. Predictions by PERCEPTIVE are shown along with confirmational wet-lab observations of DNA methylation (5mC, 5hmC, 6mA) in *N. gaditana* and 5mC in *P. soleocismus.* Best = 1, Excellent = 0.875, Better = 0.75, Good = 0.625, Fair = 0.5, Poor = 0.375, Negligable = 0.25 and Zero = 0.

To validate the prediction of CMEs capable of conferring histone modifications, we analyzed the genomes of *N. gaditana* and *T. striata,* and performed bottom-up mass-spectroscopy analysis of histones to measure levels of histone post-translational modifications for both species [32]. In this analysis, the ratio of unmodified peptides is compared to the ratio of modified peptides. For example, in *N. gaditana* for H4K20me ∼10% of peptides were unmethylated, ∼16% were mono-methylated, ∼70% were di-methylated, and the remaining 4% were tri-methylated (Figure 4A). As the CME which confers H4K20 is processive, ie., it confers mono, di, and tri methylation, all mass-spectroscopy results for H4K20me are considered positive evidence for the modification. To counter the possibility that a specific modification might be absent on account of physiological conditions, or cell cycle state, for example histone acetylation which is reduced globally in response to carbon stress in budding yeast, we measured histone modifications under varied physiological conditions, including light and dark cycle for *N. gaditana* and nitrogen starvation for *T. striata* [46]. The highest level of modified histones for any physiological condition, averaged across biological replicates, are presented in Figure 4. Overall, mass-spectroscopy analysis of *N. gaditana* cells experiencing light stress (light and dark cycle) shows significant reproducible evidence for histone acetylation and methylation, and supports the accuracy of PERCEPTIVE (Table 2, Figure 4A). Similarly, in *T. striata* mass-spectroscopy analysis of cells experiencing nitrogen stress (replete, deplete and starved) confirms the predictions by PERCEPTIVE, showing extensive histone acetylation and methylation, and supports the accuracy of PERCEPTIVE (Table 3, Figure 4B).

**Figure 4.**
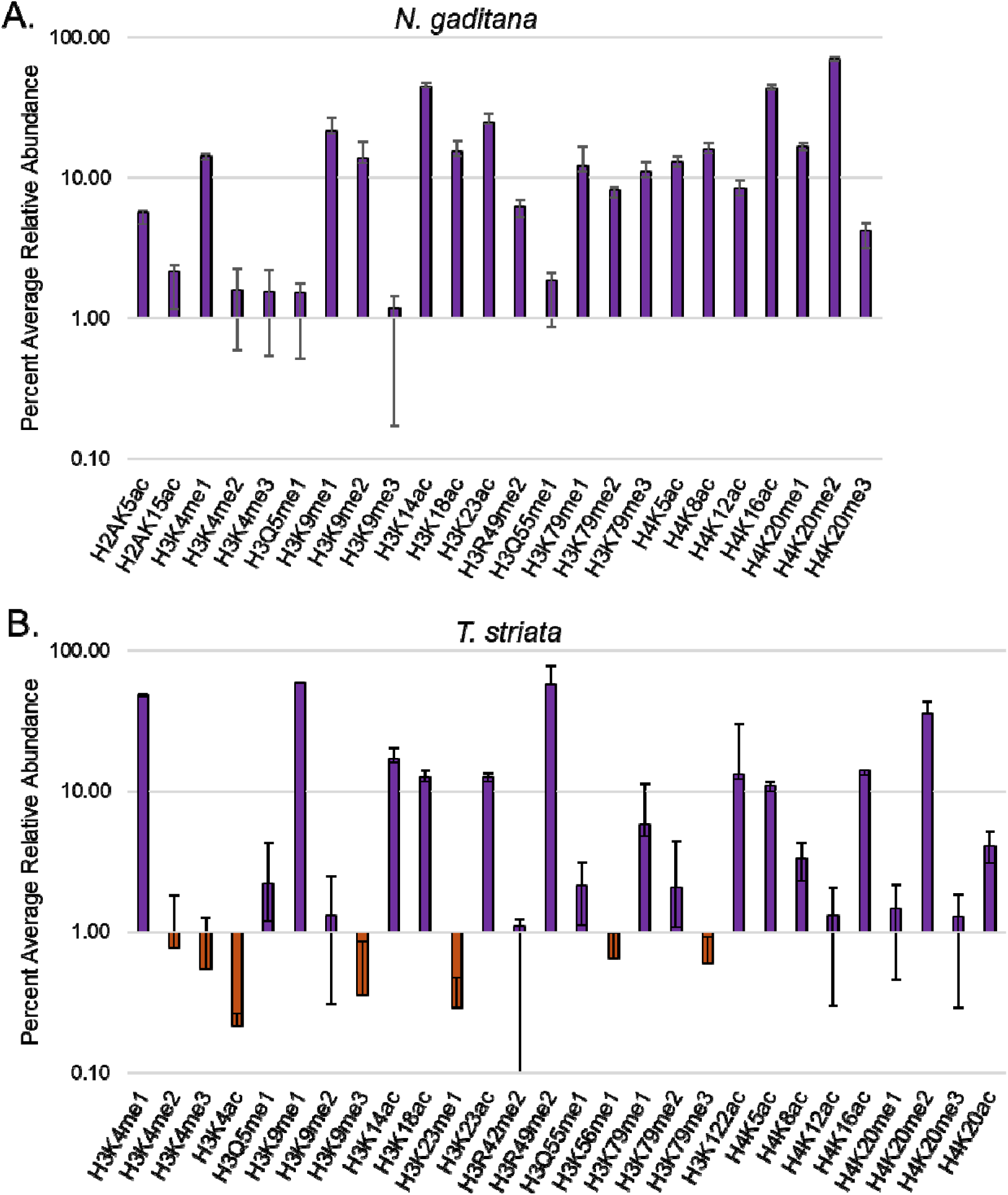
Mass spectroscopy validation of Histone Modifying Enzymes in *N. gaditana* and *T. striata.* A) Maximum percent relative abundance of various histone modifications in *N. gaditana*, measured in three biological replicates across two physiological conditions (Light/Dark). Error bars represent the standard deviation. Purple bars represent a positive finding above the experimental margin of error. B) Maximum percent relative abundance of various histone modifications in *T. striata*, measured in two biological replicates across three degrees of nitrogen stress (replete, deplete, and starvation). Error bars represent the standard deviation. Purple bars represent a positive finding above the experimental margin of error. Orange bars represent an observation within the experimental margin of error.

**Table 2.**
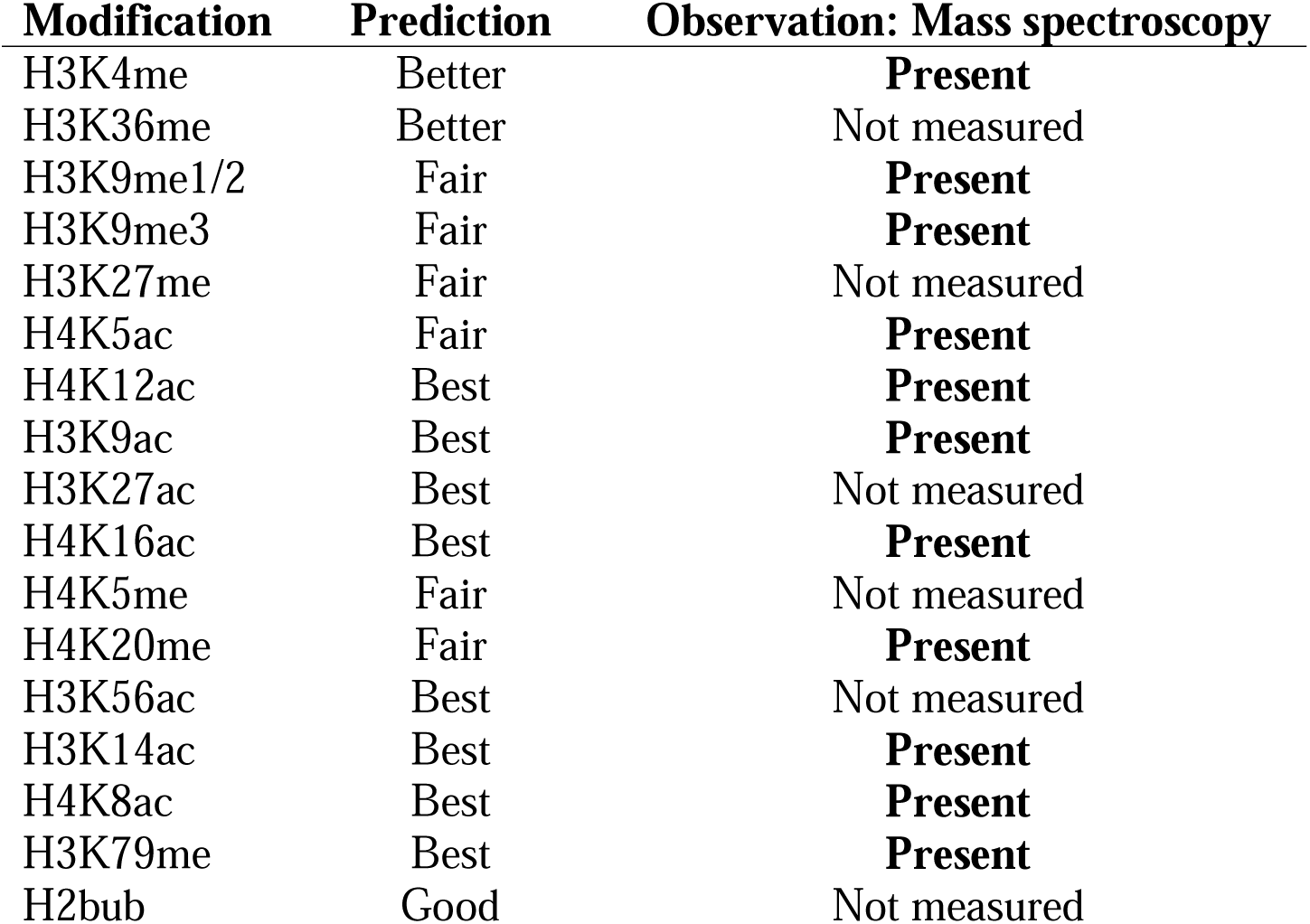
Validation of Histone Modifying Enzymes in *N. gaditana.* Predictions by PERCEPTIVE are shown along with confirmational wet-lab observations of histone PTMs in *N. gaditana*. Best = 1, Excellent = 0.875, Better = 0.75, Good = 0.625, Fair = 0.5, Poor = 0.375, Negligible = 0.25 and Zero = 0.

**Table 3.**
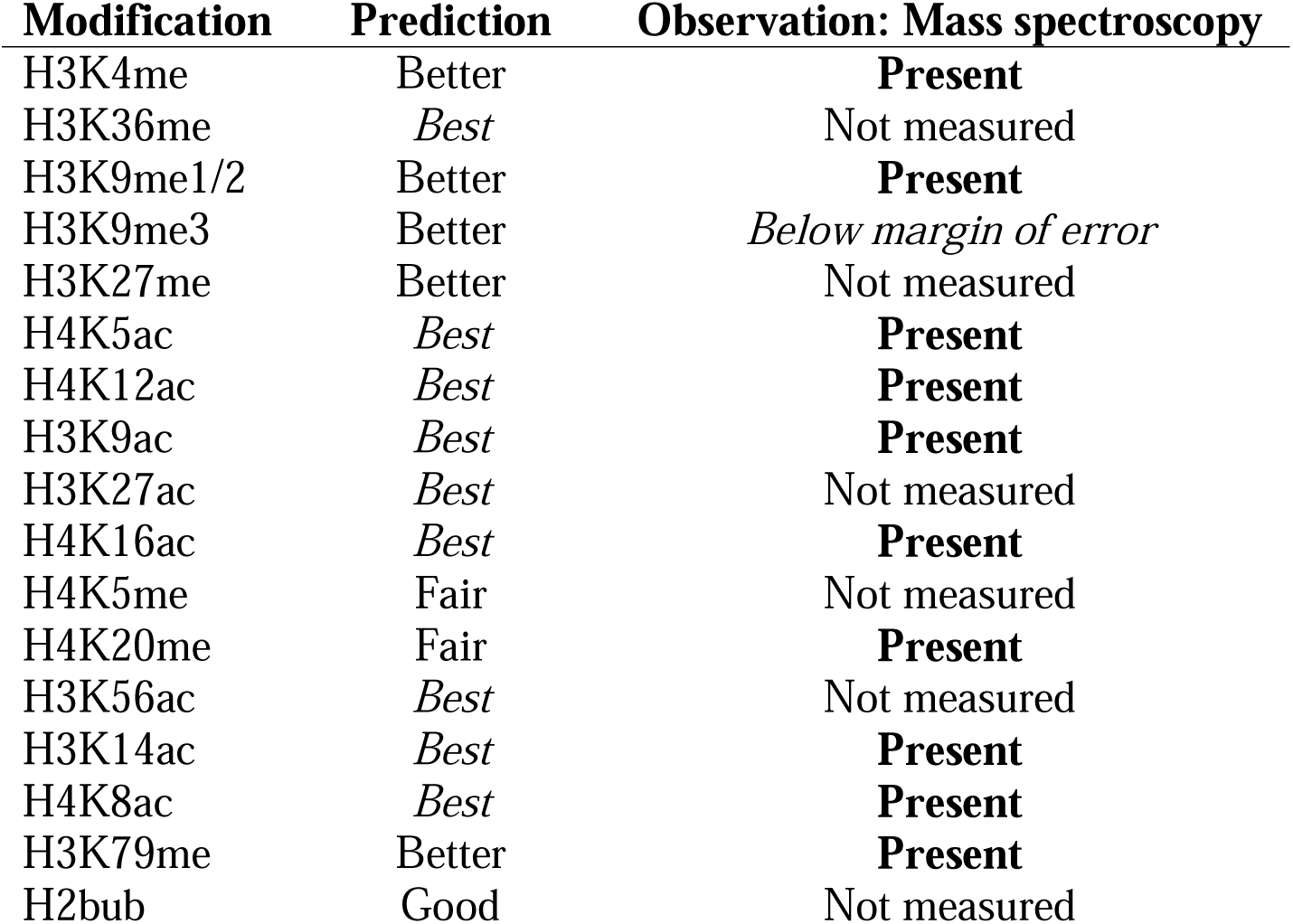
Validation of Histone Modifying Enzymes in *T. striata.* Predictions by PERCEPTIVE are shown along with confirmational wet-lab observations of histone PTMs in *T. striata*. Similarly, in *T. striata* mass-spectroscopy analysis of cells experiencing nitrogen stress (replete, deplete and starved) confirms the predictions by PERCEPTIVE, showing extensive histone acetylation and methylation, and supports the accuracy of PERCEPTIVE (Table 3, Figure 4B).

### 3.4 Antibody Selection

The vast majority of biochemical methods to measure DNA methylation and histone modifications in a semi-rapid fashion rely on antibodies mounted against an epitope of interest for identification. While this does not present a significant barrier to the identification of DNA methylation (with the exception of the poor sensitivity of many DNA methylation antibodies we have previously reported), the peptide-based antibody nature of most histone modification antibodies does [15]. The vast majority of commercially available antibodies are mounted against short 5-15mer peptide sequences of the histone N and C-terminal tails [47]. As such, sequence divergent from the histone sequence of the species for which an antibody was mounted, particularly in the largely unstructured tails of histones, can significantly interfere with the sensitivity of commercially available histone PTM antibodies when applied to a novel species in a wet-lab application. To address this issue, we have included a histone similarity/ antibody selection tool in PERCEPTIVE. This tool compares the histone sequences of human and budding yeast to the histone sequences identified for an organism of interest by HMMs specific to the core histone proteins. This graphical linear representation of the histone sequences aligns model organism sequences to predicted sequences, relying on the minimal Levenshtein edit distance, highlighting substitutions and homology (Figure 5) [43]. By utilizing this viewer, users can research appropriate commercial peptide antibodies against a PTM of interest, with the highest likelihood of performing well in a wet-lab environment. Owing to the variability and breadth of available commercial antibodies, this tool does not suggest or endorse antibodies. Users should instead research or request the peptide sequences against which an antibody has been mounted.

**Figure 5:**
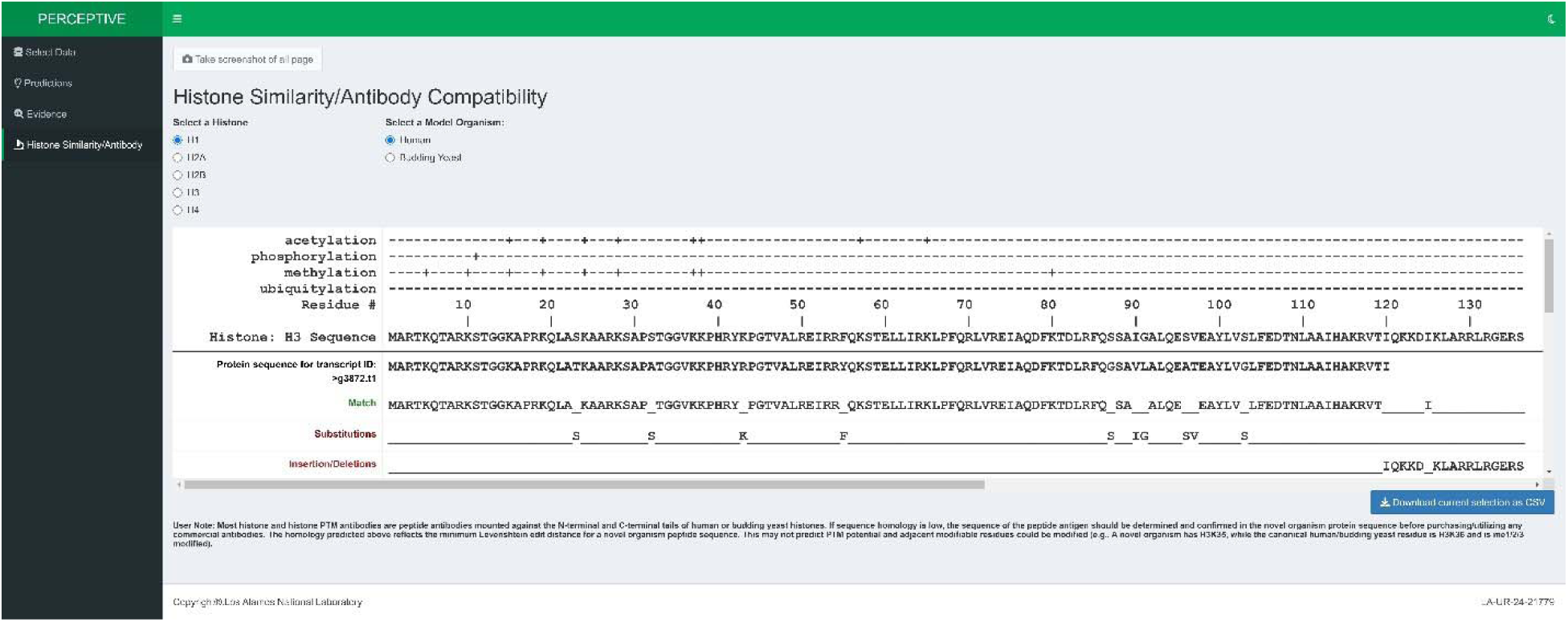
Histone Similarity and Antibody Compatibility Viewer. Linear representations of predicted histone peptide sequences are compared to sequences in model organisms with substitutions, insertions, and deletions highlighted for the user.

## DISCUSSION

PERCEPTIVE is a user-friendly graphical pipeline for the identification of chromatin modifying enzymes (CMEs) in novel species. Specifically, PERCEPTIVE provides prediction scores for DNA methylating enzymes, and histone-modifying enzymes. Moreover, the graphical interface provides users with an accessible depiction of the evidence underlying PERCEPTIVE’s predictions and informs users of best practices for next steps and wet-lab experimentation, particularly antibody selection for histone PTMs. The accuracy of our pipeline was validated against our previous observations documented in the literature for DNA methylation in *N. gaditana* and *P. soleocismus* and using histone mass spectroscopy for histone post-translational modifications (PTMs) in *N. gaditana* and *T. striata*. We believe PERCEPTIVE significantly reduces the barrier to epigenetic manipulation in non-model algae, reduces the cost and time associated with wet-lab experimentation, and focuses downstream experimental efforts.

### 4.1 Applications

Covalent modifications of DNA are most typically in the form of the addition of a methyl group to either cytosine at the 5^th^ (5-methyl-cytosine, 5mC and 5-hydroxy-methyl-cytosine, 5hmC) or 4th (4-methyl-cytosine, 4mC) position in the carbon ring, or to the 6th position in adenine (6-methyl-adenine, 6mA) [48]. While nuanced, these modifications are generally found in both eukaryotes and prokaryotes, and the addition of a methyl group is commonly associated with transcriptional regulation and sometimes repression [49]. In this case, knowing whether an organism has the potential for DNA methylation represents a strong advantage before performing costly sequencing to define the methylome, for example, the lack of 5mC predicted in *N. gaditana* by PERCEPTIVE (Table 1). However, once the methylome is defined, promoter regions near regions of endogenous hypermethylation or hypomethylation can be targeted for gene insertion and manipulation, potentially increasing output.

Comparatively, modifications of histones are more extensive and extraordinarily diverse, spanning mechanisms from transcription factor interactions at promoters to three dimensional chromatin structure [50,51]. Our focus is on the fundamental subunit of chromatin, the nucleosome core particle, which is comprised of ∼147 bp of DNA coiled around an octameric protein complex composed of 8 histone proteins [52]. Our interest in the nucleosome is in no small part due to the conservation of this complex from humans to archaea [36]. The nucleosome regulates transcription in two key ways. First, the presence of a nucleosome itself may impede the transcriptional machinery or binding of factors that promote transcription to DNA and, thereby, genes [50]. As such, cellular machinery exists to evict and place nucleosomes along DNA. Broadly, these enzymes are termed chromatin remodelers, with individual complexes having discrete roles; for example, histone chaperones facilitate the assembly of nucleosomes along DNA, while remodelers facilitate the eviction and reassembly of nucleosomes in the wake of transcription [53]. Second, nucleosomes can be covalently modified post-translationally with the addition of acetyl, methyl, phospho, ubiquitin, and other groups to the residues of the four core histone proteins (H2A, H2B, H3, and H4), which make up the nucleosome [51]. Altogether, there are at least one hundred known histone modifications, which can be placed exclusively or, in many cases, in a combinatorial fashion. These single or combinatorial modifications can promote or repress transcription by controlling the stability of the nucleosome complex, resisting or promoting chromatin remodeling, and directly signaling for or inhibiting the transcriptional machinery. For example, acetylation of histone H4 at lysine 16 (H4K16ac) promotes the relaxation of chromatin into a more transcriptionally permissive state termed euchromatin (the inverse of heterochromatin) [40,54]. Conversely, phosphorylation of histone H3 at serine 11 (H3S10p) promotes tightening of chromatin and the formation of the rigidly packaged DNA seen during mitosis, which in most species is transcriptionally repressive [55,56].

While the specific applications for the manipulation of histone PTMs are less clear, as they are more dependent on the repertoire of CMEs and PTMs in a given organism, they are equally powerful. For example, knowing that an organism has H3K79me, which acts as a barrier to the spreading of the telomeric silencing machinery, might mean that sub-telomeric regions are still transcriptionally active and that genes in these regions are less likely to be suppressed transcriptionally [57]. In another case, if an organism has H3K4me3, which is pro-transcription and marks active genes, strategies to induce targeted H3K4me3 at specific genes are likely to be interpreted positively by the cellular machinery [58]. Moreover, while not a core function of PERCEPTIVE, evidence for chromatin remodelers and chromatin structural proteins included in PERCEPTIVE may suggest that chromatin structure experiments (ATAC-seq or Hi-C), which measure chromatin accessibility and contacts, respectively, may be informative [59,60]. For example, ATAC-seq experimentation may uncover regions of chromatin that are more likely to be accessible to the transcriptional machinery, making them ideal locations to target for synthetic construct insertion.

### 4.2 Assumptions and caveats

Despite the accuracy of PERCEPTIVE achieved in our test cases, several caveats must be acknowledged. First, PERCEPTIVE cannot identify putative CMEs that lack homology and conservation of core domains found in annotated CMEs in model species. While unlikely, particularly given the breadth of organisms integrated into the InterPro database, PERCEPTIVE may underpredict a chromatin modification, for example, if a functional novel non-homologous protein, which lacks conserved domains has arisen in an organism of interest. Conversely, just because an organism has a protein with homology and conserved domains to one which confers a modification in model species does not mean that the protein is active in a novel species. Generally, while PERCEPTIVE should accurately predict most chromatin modifications, it is not a substitute for subsequent experimental work but should be used as a tool to inform downstream experimental design.

In addition to these caveats, it is important to note that while the default settings implemented in PERCEPTIVE for Canu and VELVET should provide acceptable genomes for downstream analysis, these settings assume high read quality, appropriate sequencing depth, and genomes of average eukaryotic size. Moreover, genome assembly is memory and compute-intensive and may be better done in a high-performance computing environment, where users can fine-tune assembly *prior* to PERCEPTIVE annotation. These asides underlie our suggestion that users provide assembled genomes to PERCEPTIVE and only utilize the assembly functionality if the above requirements are met with a high degree of confidence.

### 4.1 Summary

Epigenetic mechanisms are central to the regulation of organismal behavior and response to environmental perturbations. While many external factors are likely at play, the heterogeneity observed in large-scale algal cultures is at least in part likely epigenetic in nature [61]. Moreover, instances of “genetic drift” in bioengineered species are also just as likely to be epigenetic in nature. As such, we believe that any species selected as a feedstock would benefit from PERCEPTIVE analysis, even if the end goal is not epigenetic manipulation. Simply knowing that an organism may possess epigenetic plasticity could explain variable performance and inform better practices for culture and engineering. Moreover, PERCEPTIVE defines actionable first steps for epigenetic experimentation, limiting potentially fruitless work and increasing the odds of successful epigenetic engineering and manipulation. Overall, PERCEPTIVE is a powerful tool for the prediction of chromatin modification in novel organisms, with broad applications within the algae field and potentially beyond.

## PACKAGE AVAILABILITY

PERCEPTIVE is available for download via github: https://github.com/ericmsmall/PERCEPTIVE. Additionally, annotated example data for *T. striata* and *N. gaditana* are available for download.

## LICENSURE

PERCEPTIVE is an open-source package under BSD-3 copyright distribution, *Triad National Security, LLC. All rights reserved.* PERCEPTIVE utilizes the GeneMark software package from Georgia Institute of Technology within the BRAKER3 pipeline. The software is free for all academic, non-profit, and government agencies, and requires a license. For commercial applications permission must be obtained.

## AUTHOR CONTRIBUTIONS

CRS and EMS conceived the work. CRS and EMS conceived the pipeline and interface. EMS build, implemented, coded and distributed the pipeline and interface. EMS designed and performed the mass spectroscopy experiments alongside the Northwestern Proteomics Core. EMS wrote and edited the manuscript, and CRS critically revised and edited the manuscript. CRS obtained and provided funding. All authors approved the manuscript in its final form.

## ACKNOWLEDGMENTS

We would like to thank Dr. Raul Gonzalez and Sarah Pacheco for their help in providing *N. gaditana.* Additionally, we would like to thank Dr. Michael Caldwell and the members of the Northwestern Proteomics Core for their support in the mass spectroscopy component of this project.

